# Hyperexcitability precedes CA3 hippocampal neurodegeneration in a dox-regulatable TDP-43 mouse model of ALS-FTD

**DOI:** 10.1101/2024.09.24.612703

**Authors:** William Rodemer, Irene Ra, Elizabeth Jia, Jaskeerat Gujral, Bin Zhang, Kevt’her Hoxha, Bo Xing, Sanya Mehta, Madona Farag, Silvia Porta, Frances E. Jensen, Delia M. Talos, Virginia M.-Y. Lee

## Abstract

Neuronal hyperexcitability is a hallmark of amyotrophic lateral sclerosis (ALS) but its relationship with the TDP-43 aggregates that comprise the predominant pathology in over 90% of ALS cases remains unclear. Emerging evidence in tissue and slice culture models indicate that TDP-43 pathology induces neuronal hyperexcitability suggesting it may be responsible for the excitotoxicity long believed to be a major driver of ALS neuron death. Here, we characterized hyperexcitability and neurodegeneration in the hippocampus of doxycycline-regulatable rNLS8 mice (NEFH-tTA x tetO-hTDP-43ΔNLS), followed by treatment with AAV encoded DREADDs and anti-seizure medications to measure the effect on behavioral function and neurodegeneration. We found that approximately half of the CA3 neurons in the dorsal hippocampus are lost between 4 and 6 weeks after TDP-43ΔNLS induction. Neurodegeneration was preceded by selective hyperexcitability in the mossy fiber – CA3 circuit, leading us to hypothesize that glutamate excitotoxicity may be a significant contributor to neurodegeneration in this model. Interestingly, hippocampal injection of AAV encoded inhibitory DREADDs (hM4Di) and daily activation with CNO ligand rescued anxiety deficits on elevated zero maze (EZM) but did not reduce neurodegeneration. Therapeutic doses of the anti-seizure medications, valproic acid and levetiracetam, did not improve behavior or prevent neurodegeneration. These results highlight the complexity of TDP-43 - induced alterations to neuronal excitability and suggest that whereas targeting hyperexcitability can meliorate some behavioral deficits, it may not be sufficient to halt or slow neurodegeneration in TDP-43-related proteinopathies.

**Significance Statement:** Cytoplasmic aggregates of TAR DNA Binding Protein 43 (TDP-43) are the predominant pathology in over 90% of Amyotrophic lateral sclerosis (ALS) and the majority of frontotemporal lobar degeneration (FTLD-TDP) cases. Understanding how TDP-43 pathology promotes neurodegeneration may lead to therapeutic strategies to slow disease progression in humans. Recent reports in mouse and cell culture models suggest loss-of-normal TDP-43 function may drive neuronal hyperexcitability, a key physiological hallmark of ALS and possible contributor to neurodegeneration. In this study, we identified region-specific hyperexcitability that precedes neurodegeneration in the inducible rNLS8 TDP-43 mouse model. Suppressing hyperexcitability with chemogenetics improved behavioral function but did not reduce hippocampal neuron loss. Anti-seizure medications had no beneficial effects suggesting directly targeting hyperexcitability may not be therapeutically effective.

## INTRODUCTION

Neuronal hyperexcitability is a physiological hallmark of amyotrophic lateral sclerosis (ALS) yet its relationship to the underlying TAR DNA Binding Protein 43 (TDP-43) pathology and neurodegeneration remain poorly understood. Electrophysiological studies using transcranial magnetic stimulation (TMS) have consistently identified neuronal hyperexcitability both in the brain (Vucic et al. 2011) and in the periphery (Vucic and Kiernan 2006a) of ALS patient cohorts. Cortical TMS studies have shown ALS patients to present with increased motor evoked potential amplitude, a reduced cortical silent period, increased intracortical facilitation, and reduced short-interval intracortical inhibition (Vucic et al. 2011). Hyperexcitability appears early in disease (Vucic and Kiernan 2006b) and correlates with cognitive impairment (Agarwal et al. 2021; Higashihara et al. 2021). Indeed, cortical hyperexcitability may be a key distinguishing feature of ALS vs mimicking neuromuscular disorders (Vucic et al. 2011; Menon et al. 2015). Early hypothesis implicated glutamate excitotoxicity as a proximate cause of neuron death (Rothstein, Martin, and Kuncl 1992; Rothstein 1995; Roisen et al. 1982; Plaitakis and Caroscio 1987) and the anti-glutamatergic agent riluzole became the first approved ALS therapeutic (Lacomblez et al. 1996). Excitotoxicity remains central to the so called, “Dying Forward” hypothesis, which posits that excitotoxic input from upper motor neurons in layer V promotes cell death of lower motor neurons in the spinal cord (Odierna et al. 2024).

TDP-43 is predominantly nuclear, while phosphorylated cytoplasmic aggregates, with accompanying nuclear depletion, are the primary pathology found in over 90% of ALS cases (Neumann et al. 2006). Toxicity resulting from TDP-43 pathology is believed to arise from a combination of gain-of-toxic function and loss-of-normal function effects (Chen-Plotkin, Lee, and Trojanowski 2010). Recent reports in TDP-43 mouse and cell models suggest TDP-43 dysfunction induces hyperexcitability. Patch-clamp experiments in acute slice culture have demonstrated intrinsic increases in excitability among layer V pyramidal neurons in the mutant TDP-43 (Q331K) (Fogarty et al. 2016) and CamKIIα-hTDP-43ΔNLS (Dyer et al. 2021) mice. Additionally, in separate studies, layer V motor neurons from both the CamKIIa-hTDP-43ΔNLS (Dyer, Woodhouse, and Blizzard 2021) and TDP-43 (A315T) (Handley et al. 2017) mice experience widespread loss of dendritic spines consistent with sustained hyperactivity (Jiang et al. 1998).

Although evidence indicates TDP-43 dysfunction can alter neuronal activity, these alterations are complex, and it is unclear whether they lead directly to neurodegeneration. In a 2013 paper, astrocyte-specific expression of mutant TDP-43 (M337V) lead to downregulation of the glutamate transporter, GLT-1/EAAT2, that in-turn promoted excitotoxicity in neurons (Tong et al. 2013). Additionally, in CamKIIa-hTDP-43ΔNLS mice, it has been reported that layer V motor neuron hyperexcitability precedes synaptic remodeling and cell death of lower motor neurons (Reale et al. 2023). However, those changes were accompanied by potentially compensatory increases in signaling from inhibitory circuits. Additionally, patch-clamp in those same layer V neurons showed decreases in excitatory output, even as intrinsic excitability increased (Dyer et al. 2023). Thus, while multiple reports suggest TDP-43 can induce hyperexcitability, it may not necessarily drive excitotoxic neurodegeneration.

To address more directly whether hyperexcitability promotes neurodegeneration in TDP-43 proteinopathy, we characterized neurodegeneration and hyperexcitability in the hippocampus of dox-regulatable rNLS8 mice, which express mutant human TDP-43 (hTDP-43ΔNLS) in neurons throughout the brain and spinal cord, with multielectrode array electrophysiology (MEA). We then targeted hyperexcitability directly with AAV-encoded chemogenetics and anti-seizure medication (ASMs). Notably, we identified profound, selective, degeneration of the neurons in CA3 subfield of the dorsal hippocampus preceded by robust CA3 regional hyperexcitability. Interestingly, we observed a rescue of deficits on the elevated zero maze (EZM) with AAV encoded inhibitory Designer Receptors Exclusively Activated by Designer Drugs (DREADDs). No reduction in neurodegeneration was observed with treatment with either ASMs or chemogenetics.

## MATERIALS AND METHODS

### Animal Husbandry and Breeding

rNLS8 bigenic mice were generated by breeding tet-O-TDP-43ΔNLS and tTA-NEFH transgenic mice as previously described (Walker et al. 2015) and maintained on doxycycline containing chow (“on-DOX”; 200mg/kg, Bio-Serv) until approximately 2 – 6 months of age, when TDP-43ΔNLS expression was induced by replacing dox-feed with standard rodent chow (“off-DOX”). Their diet was supplemented with moistened chow pellets and DietGel76A (ClearH2O) as the motor phenotype progressed. Non-bigenic littermates and on-DOX rNLS8 were used as controls, as indicated. All animal procedures were conducted with approval by the University of Pennsylvania Institutional Animal Care and Use Committee. Approximately equal number males and females were used for all experiments.

### Accelerating Rotarod

To test motor coordination, mice were transferred to rotating arms of an accelerating rotarod (Model 7650, Ugo Basile) for 3 runs. (1) Habituation: 300s at constant 4 rpm, mice that fell during this phase were returned to the rotarod. (2, 3) Timed Trial: latency to fall (s) was assessed during 4 – 40 rpm acceleration over 300s. Mice that did not fall were assigned a latency to fall of 300s. Mice were given 3 min rest between each trial. The longest latency to fall from the 2 timed trials was used for analysis.

### Wire hang

In some studies, wire hang was used as a complement to accelerating rotarod as a general measure of strength and motor function. Mice were transferred to a wire cage lid and suspended upside down above a clean cage (approximately 30 to 60 cm). The longest latency to fall from two trials was used for analysis. Mice that remained suspended for the maximum time of 180 seconds during the first trial were not retested.

### Open field test

Gross motor function and exploratory behavior were assessed with an open field test. Individual mice were transferred to an arena (40 x 40 cm) for 15 min of free exploration (Gould, et al. 2009). Movement (distance traveled, meters) was automatically tracked by overhead camera controlled by Ethovision software (Noldus).

### Elevated zero maze

Fear and anxiety behaviors were examined with the elevated zero maze, which consists of an elevated circular platform with two walled (enclosed) quadrants on opposite sides adjacent to two open quadrants (Shepherd, et al. 1994). Mice were transferred to an enclosed quadrant for 5 min of free exploration while a video camera recorded above. Transitions were recorded when both the front and hindlimbs crossed into a new quadrant. Percent time in open quadrants was defined as the total number of seconds in the open quadrants divided by the total test time (300s).

### Y maze

Spatial working memory was assessed with the Y maze, an apparatus consisting of three walled arms (“A”, “B”, “C”) that form a “Y” shape (Kraeuter Ak, Guest PC, and Sarnyai Z. Methods Mol Biol. 2019). Mice were transferred to the distal portion of an arm, facing the center and allowed to explore for 5 min while behavior was recorded from an overhead camera. Both front and hindlimbs needed to cross the arm boundary for an arm entry to be recorded. Alternations were recorded when the mouse entered all three arms (e.g. “ABC”, “ACB”, “BCA”, “BAC”, “CAB”, “CBA”; % Spontaneous Alternations = Number of Alternations / (Number of Arm Entries – 2) x 100).

### Social interaction test

Mice were transferred to the center-chamber of a three-chambered apparatus while recorded by a video camera from above. Before the start of testing, identical plexiglass cylinders with numerous holes for air exchange were placed into the center of each of the end-chambers. During habituation (Phase I, 10 min), mice explored the apparatus with empty cylinders. In the social choice phase (Phase II, 10 min), one plexiglass cylinder was loaded with a novel same-sex stimulus mouse, while the plexiglass cylinder in the opposite end-chamber was loaded an inanimate object. For the final, direct interaction phase (Phase III, 5 min), the plexiglass cylinders were removed, allowing free interaction between the mice. Sniffing time was defined as the number of seconds the test mouse sniffed the plexiglass cylinders containing the stimulus mouse (Phase II) or sniffed / groomed the stimulus mouse directly (Phase 3). Only interactions initiated by the test mouse were scored.

### Intracerebral AAV Injection

Adult mice were anaesthetized with an intraperitoneal (i.p.) injection of ketamine (60-100 mg/kg) – xylazine (8-12 mg/kg) – acepromazine (0.5-2 mg/kg) and immobilized on a robotic stereotaxic frame (Neurostar). AAV9 encoding hM4D(Gi)-mCherry (Addgene, 50475) or mCherry control (Addgene, 114472) under the human synapsin 1 promoter (hSyn) was injected into the unilateral hippocampus (-1.94 mm anterior – posterior, +1.80 mm medial – lateral, -2.14 mm dorsal – ventral) with a 33-gauge Hamilton Syringe (2.5 µL volume, approx. 1.15 x 10^10^ genome copies total). After AAV injection, the surgical area was closed with sutures and mice monitored for recovery.

### DREADD agonist eyedrop administration

Two weeks after AAV injection, mice were taken off-DOX and began receiving daily administration of the DREADD agonist, clozapine-N-oxide (CNO) dihydrochloride (Hello Bio) for 6 weeks. CNO was diluted in sterile water to 20 mM and applied via bilateral eyedrops, using a P10 pipette, to achieve a total dose of 1mg/kg (approx. 1-2 µL per eye). Eyedrops were administered by touching eye with a droplet formed at the end of the pipette tip. The pipette tip itself did not contact the eye. If the droplet was dislodged before reaching the eye, the eyedrop was reapplied. Mice were weighed weekly to maintain proper dosage.

### Valproic acid administration

As a pharmacological alternative to the DREADDs, a subgroup of rNLS8 mice were taken off of doxycycline-chow (off-DOX) and simultaneously administered valproic acid in their drinking water for a duration of 6 weeks. Valproic acid sodium salt (Hello Bio, HB0867) was diluted in drinking water (∼1-2 mg/mL) to achieve an effective dose of approximately 300 mg kg^-1^ (mean weight of the mice in the cage (kg) x 300 mg kg^-1^ / assumed 5 mL daily water consumption). Mice were weighed weekly to maintain the appropriate dose, the drinking water – valproic solution was freshly prepared daily. Daily dosage was selected based on a previous study to achieve effective anti-convulsant activity (Ohdo, Nakano, and Ogawa 1989).

### Levetiracetam administration

rNLS8 mice were administered approx. 100 mg/kg/day levetiracetam (Selleck Chemical, S1356) in 0.5% methylcellulose and 0.1% Tween-80 by gavage (200 µL males; 130-150 µL females; calculated from the mean weight of males, and females) from 1 to 6 weeks after transgene induction (off-DOX). Gavage solutions were prepared weekly from a 400 mg/mL levetiracetam stock solution (stored in aliquots at -20°C). Control animals were administered with an equal volume of gavage vehicle alone. Daily dosage was based on a previous report to achieve anti-convulsant activity (Smucny, Stevens, and Tregellas 2015).

### Mouse tissue collection

Mice were lethally anaesthetized with a cocktail of ketamine – xylazine – acepromazine and transcardially perfused with PBS and 10% neutral buffered formalin (NBF) or PBS only (for Levetiracetam study). Brains were immersion fixed in 10% NBF overnight at 4°C then rinsed in Tris Leaching Buffer (50 mM Tris, 150 mM NaCl, pH 8.0). Post-fixed brains were grossly sectioned in the coronal plane with a mouse brain matrix and processed for paraffin embedding. Paraffin-embedded tissue blocks were cut on a rotary microtome into 6 µm thick sections, mounted on StarFrost adhesive slides, and stored at room temperature until use.

### Immunohistochemistry

Formalin-fixed paraffin-embedded sections were deparaffinized in xylene, rehydrated in graded alcohols and washed in 0.1 M Tris pH 7.6 buffer. Endogenous peroxides were quenched by incubating section in 5% H_2_O_2_ in methanol for 30 min at room temperature and antigen retrieval was performed by microwaving in citrate buffer (95°C for 15 min, room temperature for 50 min). Sections were blocked in 2% fetal bovine serum (FBS) in 0.1 M Tris buffer for 5 min then incubated with primary antibody overnight at 4°C. On Day 2, sections were incubated at room temperature with biotinylated secondary antibody for 2 hr, avidin and biotinylated horse radish peroxidase reagent (Vector Labs) for 1 hr, and ImmPACT DAB (Vector Labs) for 5-10 min. Finally, sections were dehydrated in graded alcohols, cleared with xylene, and coverglass mounted with Cytoseal 60. For immunofluorescence, peroxidase quenching was excluded, and sections were mounted with Fluoromount-G (Southern Biotech) after incubation with a fluorescently tagged secondary antibody. Primary antibodies used in this study include: rabbit anti-NeuN (ABN78, Millipore) 1:1000, mouse anti-NeuN (A60, Millipore) 1:1000, rat anti-p(409/410)-TDP-43 (clone 1D3, CNDR) 1:300, rat anti-GFAP (clone 2.2.B10, CNDR) 1:2000, rabbit anti-mCherry (26765-1-AP, ProteinTech) 1:1000, and rabbit anti-Iba1 (Wako, 019-19741) 1:1000.

### Brain slice preparation

Adult mice were deeply anesthetized with pentobarbital or a cocktail of ketamine, acepromazine, and xylazine and transcardially perfused with ice-cold dissection buffer (Sucrose 11 mM, MgCl_2_ 1 mM, CaCl_2_ 0.5 mM, NaHCO_3_ 26 mM, Glucose 10 mM, KCl 3 mM, NaH_2_PO_4_ 1.25 mM; 290 – 310 mmol/kg). Brains were quickly dissected in chilled oxygenated dissection buffer and one brain hemisphere was transferred to a vibratome (Leica VT100s) to generate horizontal brain sections (300 µm). Slices were incubated in oxygenated artificial cerebrospinal fluid (ACSF; MgSO_4_ 1.2 mM, CaCl_2_ 2 mM, NaHCO_3_ 26 mM, Glucose 10 mM, NaCl 124 mM, KCl 5 mM, NaH_2_PO_4_ 1.25 mM; 290 – 310 mmol/kg) for 50 min before recording (approx. 32°C for 30 min, then room temperature for 20 min).

### Extracellular recordings

A multielectrode array (MEA) recording system (MED64, Alpha Med Scientific) was used to perform evoked field excitatory postsynaptic potential recordings (fEPSP). The MEA probes had 64 planar electrodes in a 8 x 8 pattern with 150 µm interelectrode spacing (MED-PD5155) and were continuously perfused at room temperature with oxygenated ACSF containing 60 µM picrotoxin to block inhibitory GABAergic currents. Slices were transferred to the MEA chamber and the hippocampus was aligned to the electrode array. Slices were anchored with a harp and allowed to recover for 20 min before recording. Hippocampal subfields were identified by anatomical landmarks under brightfield microscopy. Stimulation and analysis channels were selected by assessing maximal fEPSP response to single electrical pulse among electrodes in the region of interest. Input-output curves were captured for the mossy fiber – CA3 and CA3 – CA1 circuits (5-100 µA, 5 µA steps, 30s interstimulus interval). In some recordings, slices needed to be repositioned after CA3 recordings to align the CA1 region to the electrode array. Traces were analyzed with custom code in R to identify max fEPSP amplitude per pulse stimulus Analysis was performed on the mean fEPSP amplitudes recorded from 1 – 3 tissue slices per mouse.

### Experimental Design and Statistical Analysis

Statistical analysis was performed with Graphpad Prism 10 using two-tailed t-Tests (α = 0.05), one-way ANOVA, two-way ANOVA, or mixed effects models with post-hoc multiple comparisons test as indicated. Figures were prepared using Adobe Photoshop (ver. 25.11).

### Code Accessibility

R code will be provided upon request.

## RESULTS

### Region-specific neurodegeneration in the hippocampus of rNLS8 mice

rNLS8 transgenic mice express full-length human TDP-43 with a defective nuclear localization signal (TDP-43ΔNLS) under the control of the neurofilament heavy subunit promoter (Walker et al. 2015). Transgene expression is induced by replacing doxycycline-chow with normal feed, resulting in cytoplasmic TDP-43 accumulation and subsequent depletion of endogenous nuclear TDP-43. Motor deficits begin around 2 weeks off-DOX, and neurodegeneration, evidenced by decreased cortical thickness and neuromuscular junction denervation begin around 4 weeks off-DOX (Walker et al. 2015). To quantify hippocampal neuronal loss over the disease course, we performed cell counts of the dorsal hippocampus in coronal sections of off-DOX rNLS8 mice **(Fig 1).** There was no hippocampal neuronal loss during the first 4 weeks off-DOX, while at 6-8 weeks approximately half of the neurons in the CA3/CA2 subfield are lost. Mild neuron loss was also detected in the dentate gyrus (DG) beginning at 8 weeks off-DOX. Interestingly, no cell loss was observed in the CA1 region, despite high expression of TDP-43ΔNLS.

**Figure 1.**
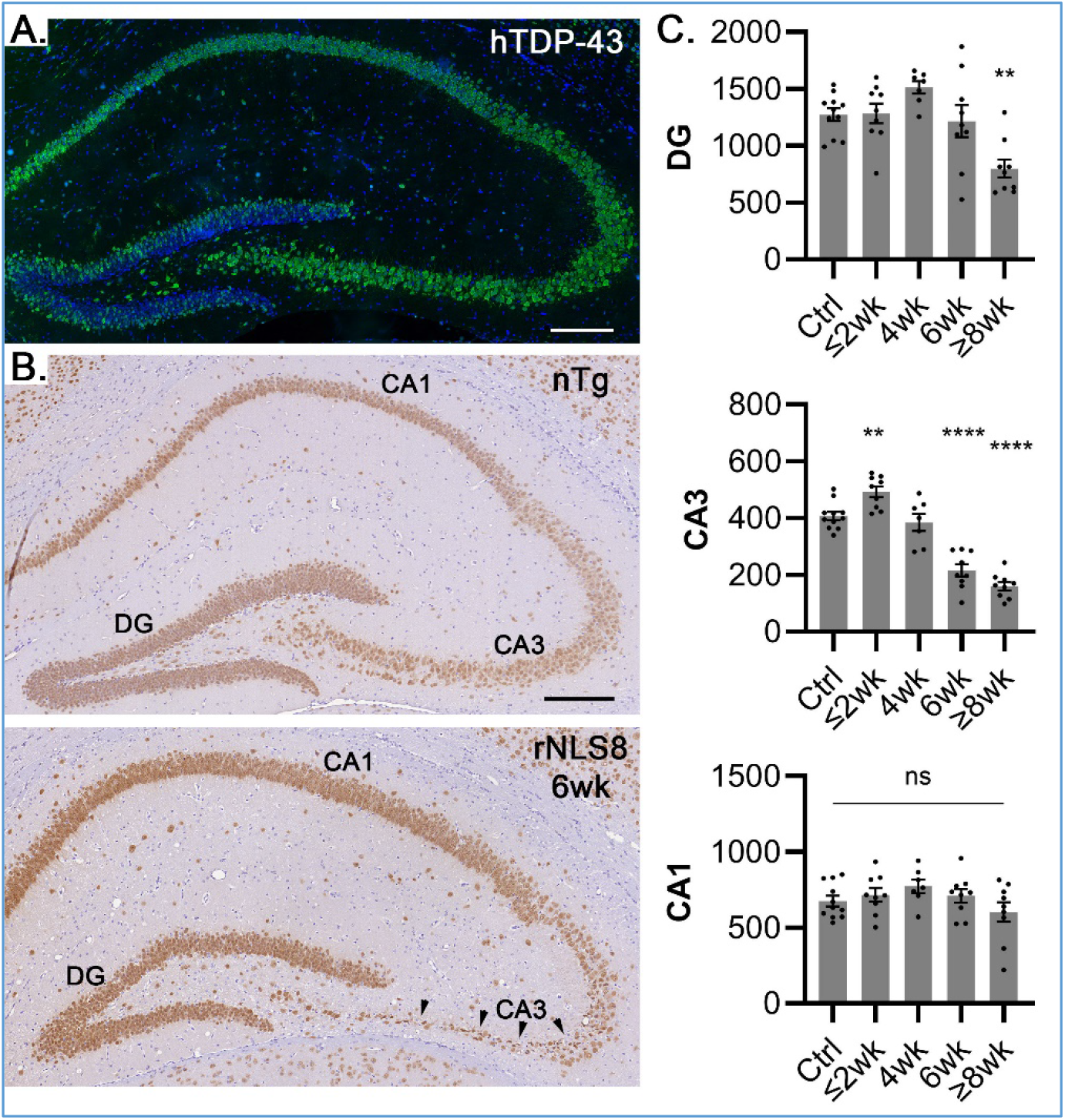
Selective death of CA3 neurons in the dorsal hippocampus between 4 and 6 weeks off-DOX in rNLS8 mice. (**A**) Immunofluorescence of cytoplasmic hTDP-43 (antibody 5104, green; DAPI, blue) in a 2 week off-DOX rNLS8 mouse (**B**) Representative NeuN immunohistochemistry in nTg control and 6 week off-DOX rNLS8 hippocampus (arrows show CA3 thinning). (**C**). NeuN(+) cell counts per section in the dentate gyrus CA3, and CA1 subfields (n = 11 control (n= 2 on-DOX rNLS8, n = 9 nTg littermates); n = 9, ≤2wk off-DOX; n = 7, 4wk off-DOX; n = 9, 6wk off-DOX; n = 9, ≥8wk off-DOX). Mean ± error bars ( S.E.M.) Scale bars, 200 µm. One-way ANOVA with Dunnett’s multiple comparisons test. **, p<0.01. ****, p<0.0001. Between 4 and 6 weeks, approximately 47-56% of dorsal hippocampus CA3 neurons are lost in off-DOX rNLS8 mice. Mild degeneration is seen in the dentate gyrus at late timepoints while CA1 appears unaffected.

### Neurodegeneration of CA3 hippocampal neurons is preceded by network hyperexcitability

Given that the DG is well-preserved in off-DOX rNLS8 mice, we hypothesized that neurodegeneration of CA3 neurons may be due to glutamate excitotoxicity from excessive excitatory input from the mossy fibers. To answer this question, we investigated hyperexcitability in the rNSL8 hippocampus using MEA electrophysiology **(Fig 2**). Here, we evoked excitatory post-synaptic potentials (fEPSPs; 0-100 µA stimulation, in 5 µA steps) from non-transgenic littermate controls and 3.1 – 4.1 week off-DOX rNLS8 mice (n = 4) in CA3 and CA1 hippocampal subfields. To limit confounding by inhibitory GABAergic circuits, all experiments were performed in the presence of 60 µM picrotoxin. Consistent with our hypothesis, we observed a dramatic increase in excitability in the mossy fiber – CA3 circuit, while the Schaffer collateral – CA1 circuit appeared unaffected. We suspected that this difference in excitability may be responsible for CA3 neuron vulnerability.

**Figure 2.**
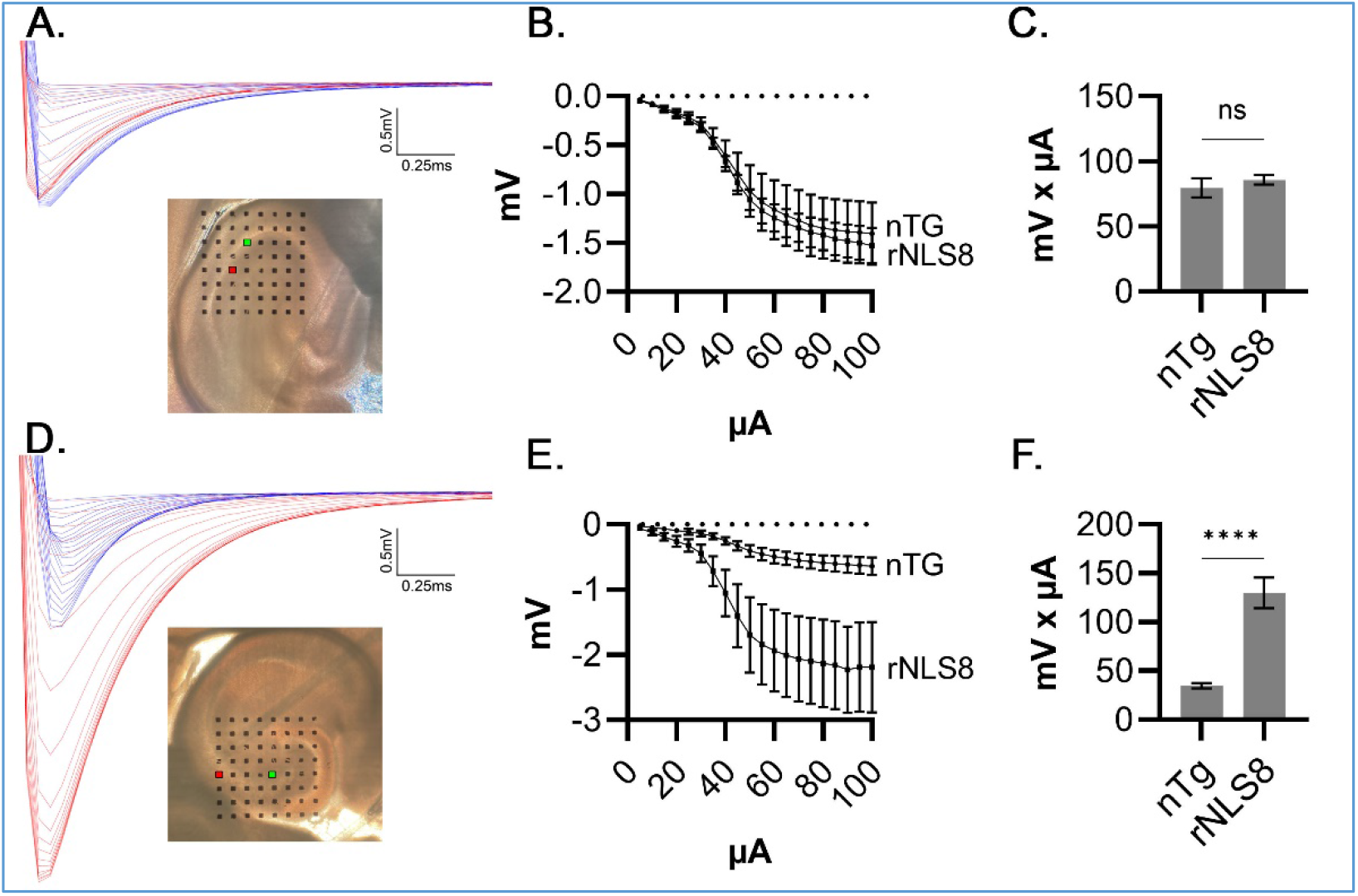
Hyperexcitability precedes neurodegeneration in CA3 hippocampus of off-DOX rNLS8 mice. **(A-C)** Evoked fEPSPs in CA1 from nTg littermates and 3.5 – 4.5 week off-DOX rNLS8 mice (n = 4 mice, 1-3 slices per mouse) showed similar amplitudes. **(D-F)** However, rNLS8 CA3 hippocampus displayed much greater excitability than nTg controls. **(A, D)** Representative fEPSP traces (nTg, blue; rNLS8, red) and micrographs of hippocampal slices (150 µm distance between electrodes; red square, stim electrode, green square, recording electrode). **(B, E)** Input-output curves of fEPSP amplititude (5-100 µA stimulation, in 5 µA steps, 30s interstim interval). **(C, F)** Area under the curve analysis. Mean ± error bars (S.E.M.). Two-tailed t-test. ****, p<0.0001.

### Cognitive and social behavioral deficits precede neurodegeneration in off-DOX rNLS8 mice

It is possible dysfunction in network excitability induces behavioral alterations prior to the onset of frank cell loss. To characterize early changes in cognitive function we subjected a cohort (n = 8) of rNLS8 mice to a weekly behavioral battery, assessing motor and cognitive function, from baseline to 4 weeks off-DOX (**Fig 3**). Off-DOX rNLS8 experienced weight loss and progressive motor impairments consistent with previous reports (Walker et al. 2015). At 4 weeks, off-DOX mice experienced a 68% reduction in performance on the rotarod assay vs on-DOX controls (75 vs 236 seconds mean latency to fall) and lost approximately 22% of their baseline weight (vs +10% weight gain for on-DOX controls).

**Figure 3.**
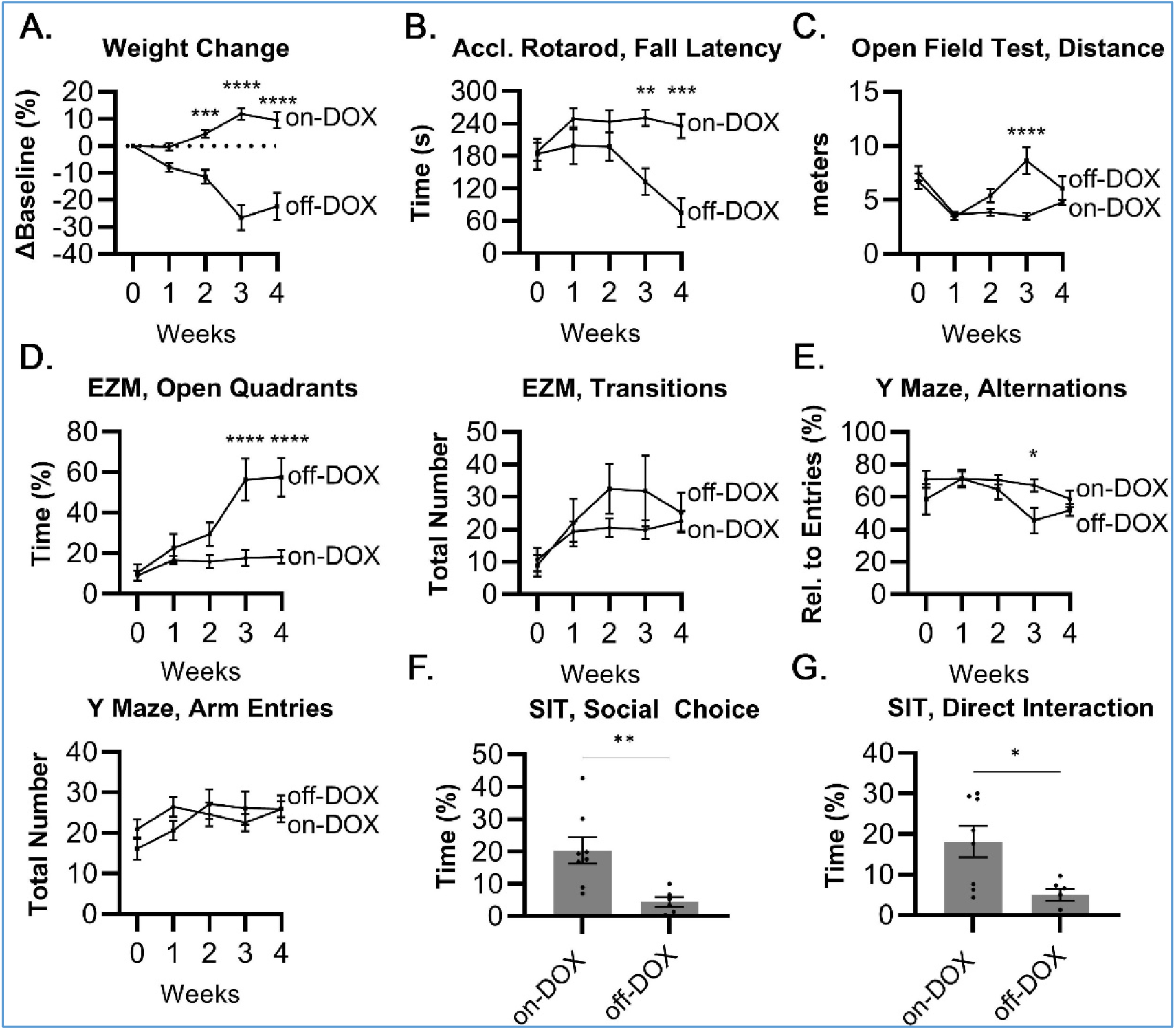
Early motor and cognitive deficits in off-DOX rNLS8 mice. **(A)** Weight change from on-DOX baseline (“Week 0”). **(B)** Latency to fall in accelerating rotarod. **(C)** Total exploration distance in a 15 min open field test. rNLS8 mice experienced progressive loss of motor coordination but retained the ability to ambulate to 4 weeks off-DOX. **(D)** Percent-time in open quadrants and total number of quadrant transitions on elevated zero maze. rNLS8 mice experienced profound deficits in anxiety-, fear-behaviors beginning at 3 weeks off-DOX. (**E)** Percent-spontaneous alternations and total number of arm entries on Y maze. A mild transient deficit was observed at in spatial memory at 3 weeks off-DOX.. Impaired sociability was observed on three-chambered social interaction test at 4 weeks as assessed by time sniffing during **(F)** social choice (restrained stimulus mouse) and **(G)** direct interaction (unrestrained stimulus mouse) phases. n = 8 each on-DOX control and off-DOX rNLS8 mice at 0, 1, 2, and 3 weeks; n = 6 off-DOX rNLS8 at 4 weeks and for social interaction test. Mean ± error bars (S.E.M.) Mixed-effects model with Šidák multiple comparison’s test (A-E) and two-tailed t-test (F-G). *, p < 0.05; ***, p < 0.01; **** p < 0.0001.

Nevertheless, motor deficits did not negatively impact exploratory behavior in the open field assay, ruling out potential secondary effects on behavioral measures. Interestingly, we observed transient increase in exploratory behavior at the 3 week timepoint in off-DOX mice. Hyperlocomoter activity is a hallmark characteristic of CamKIIa-NLS4 mice (Alfieri, Pino, and Igaz 2014). These mice express the same mutant hTDP-43ΔNLS construct as the rNLS8 but under a brain-specific promoter (CamKIIa), largely sparing them the severe deficits observed in the rNLS8 mouse line. It is possible that network hyperexcitability or other supraspinal cellular dysfunction, induce a similar hyperlocomoter phenotype in rNLS8 mice that is ultimately masked by lower motor neuron dysfunction later in the disease course.

Surprisingly, we observed substantial deficits in anxiety-like behavior deficits on elevated zero maze beginning at 3 weeks off-DOX. Multiple limbic structures including amygdala and hippocampus have been implicated in mediating the elevated plus- and zero-maze behaviors (Silveira, Sandner, and Graeff 1993; File and Gonzalez 1996; Walf and Frye 2007). It is possible, that the hippocampal dysfunction we observed on MEA and later observed undergoing neurodegeneration may contribute to these deficits. However, we observed only minimal, transient, deficits on Y maze (spontaneous alterations), a more straightforward assessment of spatial memory and hippocampal function.

Sociability deficits have been previously reported in reported in CamKIIa-NLS4 (Alfieri, Pino, and Igaz 2014). To investigate whether sociability was also impaired in rNLS8 mice, we performed a three chambered social interaction at 4 weeks off-DOX. Similar to CamKIIa-NLS4 mice, off-DOX rNLS8 mice display reduced social interactions with a novel same-sex mouse.

### Suppressing excitability with AAV encoded DREADDs reduces cognitive deficits but not gross neurodegeneration in the hippocampus

To directly address whether hyperexcitability contributed to neurodegeneration in rNLS8 mice, we performed unilateral hippocampal injection of AAV encoding the inhibitory Designer Receptors Exclusively Activated by Designer Drugs (DREADDs), i.e. hM4Di-mCherry, or mCherry only (**Fig 4**) in a cohort of N=20 rNLS8 mice (with n = 5 on-DOX controls). As a previous report in macaques indicated that that AAV encoded hM4Di receptors could attenuate bicuculline -induced discharges (Miyakawa et al. 2023), we hypothesized that DREADDs could likewise attenuate TDP-43 -induced hyperexcitability. TDP-43ΔNLS expression was induced by taking the mice off-DOX, approximately 2 weeks after AAV injection. The receptors were activated daily with administration of DREADD agonist clozapine-N-oxide (CNO), 1 mg/kg) beginning at day of off-DOX and continuing for 6 weeks. As localized DREADD activity may have been insufficient to alter the disease course, we treated an additional group of rNLS8 mice (n = 9) with the ASM valproic acid (VPA), dissolved in drinking water (to achieve 300 mg/kg/day based on mean cage weight). In SOD1 (G93A) mice, VPA administration in drinking water was shown to prolong survival (Sugai et al. 2004), thus, we expected VPA to provide a therapeutic benefit in rNLS8 mice. Contrary to our hypothesis, histological analysis at 6 weeks off-DOX revealed no improvement in neuron survival in the vulnerable hippocampal CA3 field in the DREADDs or VPA treated groups.

**Figure 4.**
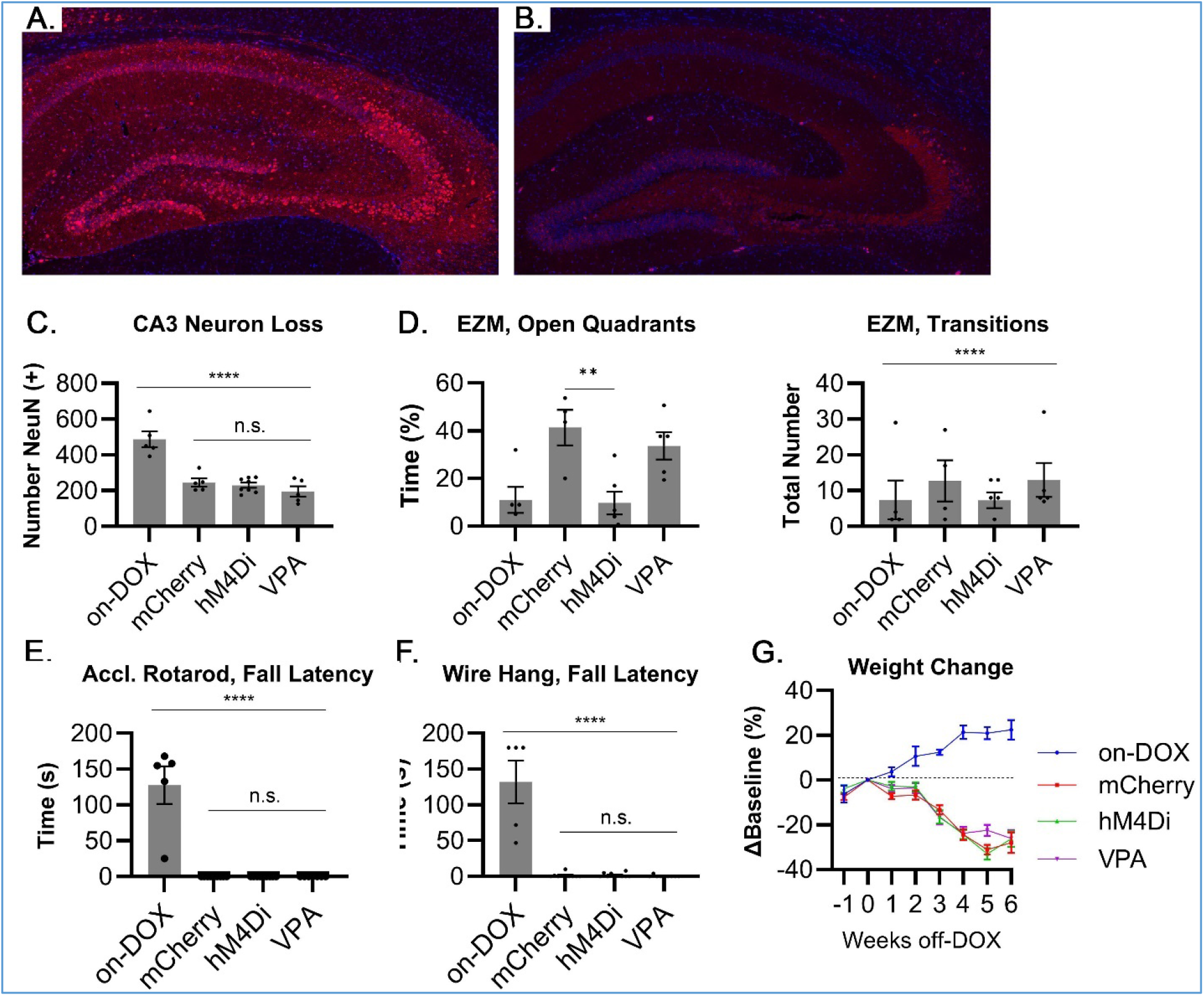
Hippocampal injected AAV-DREADDs (hM3Di-mCherry) improved fear-, anxiety-deficits without affecting neurodegeneration. Representative immunohistochemistry for mCherry in **(A)** mCherry vector controls and **(B)** inhibitory DREADD injected rNLS8 hippocampus at 6 week off-DOX (mCherry, red; DAPI, blue). At the same titer, mCherry immunofluorescence was far more intense than mCherry tagged DREADDs. Note, robust transduction of the mossy fibers in the DREADD injected mice. (**C**) No reduction in neurodegeneration was observed in counts of NeuN-positive neurons in CA3 of the dorsal hippocampus. **(D)** However, DREADD injected mice showed dramatic improvements in the Elevated zero maze. Motor performance in **(E)** accelerating rotarod or **(F)** wire hang, as well as **(G)** weight loss were similar across all off-DOX groups. Mean ± error bars (S.E.M.). One-way (C-D, F-G) and two-way (C) ANOVA with Tukey’s multiple comparisons test. **, p<0.01; ****, p<0.0001

However, we observed a dramatic rescue of anxiety-fear behaviors, prior to tissue collection at 5.8 weeks off-DOX, on the elevated zero maze in the DREADDs cohort but not in VPA treated mice. Given the dramatic behavioral rescue with DREADDs, we suspected that insufficient DREADDs activation or variability in VPA consumption may have been responsible for the lack of improvements in neuron survival.

To address this variability in dosing, we tested an ASM with a different mechanism of action than VPA. We performed a final study, where we administered, levetiracetam (LEV) daily by oral gavage (to achieve 100 mg/kg), or methylcellulose vehicle in rNLS8 mice (n=16) from 1-6 weeks off-DOX (**Fig 5**). Behavioral tests were performed at 4 weeks off-DOX and histology at 6 weeks. Previously, LEV was shown to reduce cognitive dysfunction in APP23/MAPT mice (Zheng et al. 2022) and reduce aberrant network activity in hAPPJ20 mice (Sanchez et al. 2012) when administered at comparable dosing. To enhance our ability to detect changes in the rNLS8 disease course, we included a social interaction test, where we showed previously that rNLS8 mice to have deficits **(Fig 3)**, and additional immunostaining for markers of astrogliosis and microgliosis. However, similar to VPA treatment, we observed no reduction in neurodegeneration in the CA3 subfield of the dorsal hippocampus or amelioration of behavioral deficits in LEV treated mice. Unexpectedly, we observed a mild reduction in sociability in LEV mice in the three-chambered social interaction test. Although, to our knowledge, impairments in sociability have not been shown in mice administered LEV, mood alterations including irritability, as well as fatigue are common side effects of LEV treatment in humans, which could have contributed to reduced sociability in our study.

**Figure 5.**
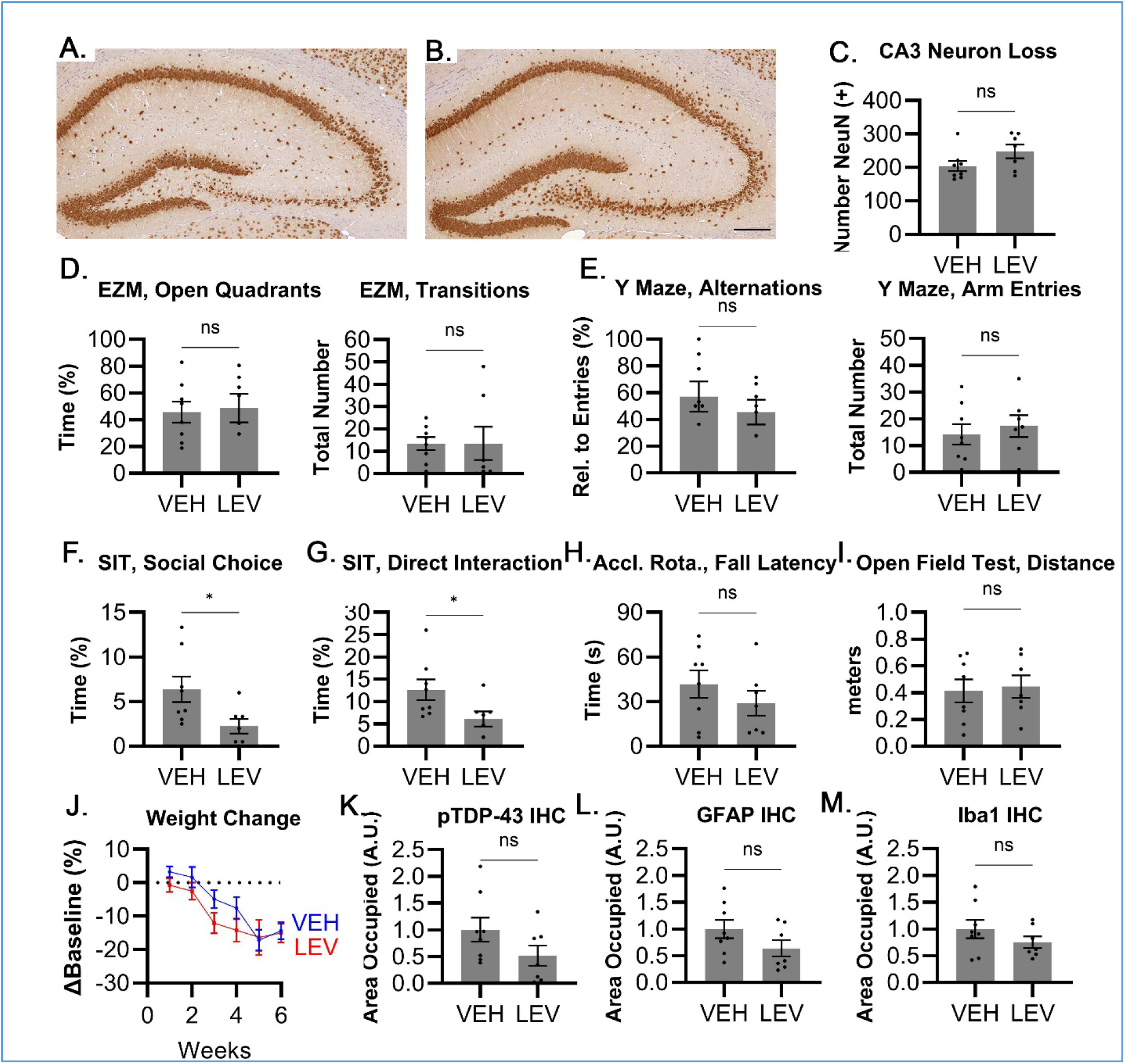
Anti-seizure medication therapy in rNLS8 mice with levetiracetam did not reduce neurodegeneration at 6 weeks off-DOX or improve behavioral deficits at 4 weeks off-DOX. Representative NeuN immunohistochemistry in 6wk off-DOX (**A**) vehicle control (VEH; n = 8) and (**B**) levetiracetam (LEV; n = 7) treated mice. (**C**) CA3 cell counts and function on the (**D**) elevated zero and (**E**) Y mazes were similar between VEH and LEV mice. Unexpectedly we observed mild reduced sociability in LEV -treated mice in both (**F**) social choice (restrained stimulus mouse) and (**G**) direct interaction (unrestrained mouse) phases. No differences between groups were observed on (**H**) accelerating rotarod, (**I**) 15 min open field test, (**J**) weight loss, or (**K**) p(409/410)TDP-43, **(L)** GFAP, or **(M)** Iba1 immunoreactivity in the dorsal hippocampus. Mean ± error bars (S.E.M.). Two-tailed t-tests (C-I, K-M) and (J) mixed effects model with Šidák multiple comparison’s test. *, p < 0.05.

## DISCUSSION

Hyperexcitability is a defining feature of ALS but its relationship to TDP-43 pathology and neurodegeneration are unclear. The molecular basis for hyperexcitability in ALS is not fully understood but likely involves a complex dysregulation of glutamatergic neurotransmission that increases susceptibility to excitotoxicity. Post-mortem studies with ALS patients have identified depletion of the astrocytic glutamate transporter (EAAT2) (Rothstein et al. 1995) in spinal cord and motor cortex, along with decreased neuronal expression of the glutamate ionotropic receptor AMPA type subunit 2 (GluA2), which regulates AMPA receptor calcium permeability, in spinal cord motoneurons (Kawahara et al. 2004). Interestingly, EAAT2 and AMPAR dysregulation have been identified in SOD-1 (Tortarolo et al. 2006; Van Damme et al. 2005; Howland et al. 2002), c9ORF72 (Selvaraj et al. 2018), FUS (Kia et al. 2018), and TDP-43 (Dyer et al. 2021; Jiang et al. 2019; Tong et al. 2013; Polymenidou et al. 2011; Wright et al. 2021) ALS experimental models, which suggests excitotoxicity may represent a common disease mechanism. Evidence that TDP-43 can alter excitability is strengthen by previous studies indicating TDP-43 regulates the RNAs encoding the AMPAR subunits: GluA2, GluA3, and GluA4 (Sephton et al. 2011). In addition to AMPAR and EAAT2, emerging literature suggests TDP-43 proteinopathy may result in dysregulation of the voltage gated potassium channel, KCNQ2 (Seddighi et al. 2024), which regulates the threshold for action potential (AP) firing and helps prevent continued AP firing after bursting (Simkin et al. 2021).

Here, we observed that rNLS8 mice experience profound neurodegeneration in the CA3 subfield of the dorsal hippocampus between 4 and 6 weeks off-DOX. Additionally, despite high cytoplasmic expression of TDP-43ΔNLS, neurons in CA1 appeared relatively unaffected. Unlike the CamKIIa-NLS4 mice, we did not observe more than mild degeneration of the dentate gyrus. As the dentate gyrus was relatively preserved, we hypothesized that excessive excitatory input from mossy fibers may promote CA3 neurodegeneration via glutamate excitotoxicity. To characterize hyperexcitability, we performed multielectrode array electrophysiology in acute hippocampal slices from off-DOX rNLS8 mice, where we observed robust hyperexcitability in CA3 but not CA1 subfields.

In CamKIIa-NLS4 mice, corticomotor neuron hyperexcitability preceded synaptic remodeling and degeneration of lower motor neurons (Dyer et al. 2021). In our study, region-specific hyperexcitability preceded selective neurodegeneration in the CA3 subfield. Thus, we hypothesized that excessive excitatory input from the dentate gyrus, induced excitotoxicity among CA3 pyramidal neurons. To test this hypothesis, we used three approaches to suppress hippocampal hyperexcitability. First, we use AAV encoded DREADDs injected locally into the hippocampus, which we activated with daily eyedrop of C21 ligand. In a separate cohort, we administered the commonly used anti-seizure medication, valproic acid, in the drinking water. Interestingly, we observed a rescue of anxiety-deficits on the elevated zero maze but no corresponding reduction in neuron loss. The valproic acid had no measurable effect. As disease progression may alter water consumption in rNLS8 mice, it is possible that the dose the mice received varied unexpectedly. To address this issue, we performed a final study where we administered the anti-seizure medication, levetiracetam, by oral gavage. Like valproic acid treatment, this had no effect on neurodegeneration or behavioral deficits.

Results from this study suggest that directly targeting excess excitability with pharmacology may not be effective therapeutic approach in TDP-43 proteinopathies. Although our behavioral rescue with the inhibitory DREADDs suggests that it may have the potential to address symptoms. This is not completely surprising as emerging reports suggest TDP-43 -induced excitability changes are highly complex and involve potential intrinsic and extrinsic compensatory mechanisms including alterations in excitatory output (Dyer et al. 2023), increased inhibitory signaling (Reale et al. 2023), and potentially novel protective effects of microglia (Xie et al. 2024) or other non-neuronal cell types. Many questions remain. Most notably this includes whether changes in excitability contribute to TDP-43 disease phenotype and how best to address those changes therapeutically. Additionally, within the rNLS8 model, CA3 neurodegeneration is striking but the mechanism remains unclear.

## Acknowledgements

“This work was supported by NIA, 5T32AG000255; NINDS, 5R01NS110688; and a seed grant from the American Epilepsy Society (AES).”

## Notes

**Conflict of interest statement:** “The authors declare no competing financial interests.”

### Competing Interest Statement

The authors have declared no competing interest.

